# Maintenance DNA methylation is necessary for age-related alterations in regulatory T cell transcriptional and DNA methylation signatures

**DOI:** 10.64898/2026.06.24.733005

**Authors:** Jonathan K Gurkan, Qianli Liu, Carla P Reyes Flores, Kathryn A Helmin, Daniel H Ryan, Anthony M Joudi, Benjamin J Ulrich, Hiam Abdala-Valencia, Elizabeth M Steinert, Benjamin D Singer

## Abstract

CD4^+^FOXP3^+^ regulatory T (Treg) cells maintain self-tolerance, restrain immune responses during inflammatory stimuli, and promote tissue function and repair. Treg cell lineage identity, stability, and function depend on specific DNA methylation patterns maintained by the epigenetic regulator, UHRF1. Aging disrupts DNA methylation patterns necessary for Treg cell-mediated lung repair in a cell-autonomous manner. Nevertheless, whether maintenance DNA methylation is necessary for age-related Treg cell transcriptional and methylation programs is unknown. Here, we performed transcriptional and DNA methylation profiling on young and old Treg cells isolated from mice with chimeric Treg cell-specific loss of UHRF1. We observed cell-autonomous, age-related alterations in transcriptional and DNA methylation signatures that were dependent on UHRF1. We conclude that maintenance DNA methylation is required for age-related alterations in Treg cell transcriptional and DNA methylation signatures.

## Introduction

Regulatory T (Treg) cells are a subset of CD4^+^ T cells that maintain immune self-tolerance, suppress overexuberant effector immune cell responses during inflammatory stimuli, and promote repair following injury to the lung and other organs^1–5^. Treg cells require a specific epigenetic pattern marked by differential cytosine-phospho-guanine (CpG) DNA methylation at specific genomic loci compared with conventional CD4^+^ T cells^6^. Aging-related epigenetic changes, particularly in DNA methylation, disrupt the function of multiple immune cell subsets, including Treg cells^7^. We showed that aging imparts a cell-autonomous defect in Treg cell reparative function by disrupting DNA methylation patterning^8^. Mechanistically, we determined that maintenance of Treg cell DNA methylation patterns by the epigenetic regulator, ubiquitin-like with plant homeodomain and RING finger domains 1 (UHRF1), is necessary for Treg cell lineage stability and function over time in young adult mice^9,10^. Mice with Treg cell-specific loss of UHRF1 developed the *scurfy* phenotype and displayed de-repression of effector T cell loci (e.g., *Tbx21*) alongside global DNA hypomethylation, leading to destabilization of Treg cell lineage commitment, spontaneous inflammation, and a secondary gain of methylation at the *Foxp3* locus. Given the importance of Treg cell function in a variety of inflammatory disease states and potential efficacy as a cellular therapy^4,11,12^, particularly in older adults, further understanding of the epigenetic determinants of Treg cell function throughout the lifespan is essential. Here, we demonstrate that UHRF1-mediated maintenance DNA methylation is required to preserve transcriptional and DNA methylation signatures in Treg cells in an age-dependent and cell-autonomous manner.

## Results

To test the necessity of UHRF1-mediated maintenance DNA methylation in determining age-related transcriptional and DNA methylation signatures in Treg cells, we bred *Uhrf1*-chimeric female mice (*Uhrf1*^*fl/fl*^*Foxp3*^*+/YFP-Cre*^)^9^. Briefly, these mice contain a mixed population of UHRF1-sufficient and UHRF1-deficient Treg cells, because random inactivation of the X chromosome prevents *Foxp3*-driven Cre-recombinase activity in roughly half of the total population of Treg cells. UHRF1-deficient Treg cells lose *Foxp3*-YFP expression over time, as they lose *Foxp3* expression in their transition to ex-FOXP3 cells^9^. Regardless, we showed that *Foxp3*-YFP^+^ Treg cells in this model are UHRF1 deficient^9^. In addition, since the UHRF1-sufficient population within each chimeric mouse is sufficient to maintain self-tolerance, this strategy allows for the analysis of cell-intrinsic differences in the absence of an inflammatory microenvironment^9^.

We hypothesized that transcriptional signatures in Treg cells depend on maintenance DNA methylation throughout the lifespan. To test this hypothesis, we performed transcriptional profiling via RNA-sequencing of splenic UHRF1-sufficient (CD3ε^+^CD4^+^*Foxp3*-YFP^−^CD25^hi^FR4^hi^)^13^ and deficient (CD3ε^+^CD4^+^*Foxp3*-YFP^+^) Treg cells isolated from young (3-month-old) and old (20-24-month-old) *Uhrf1*-chimeric female mice (**Supplemental Figure 1**).

**Figure 1.**
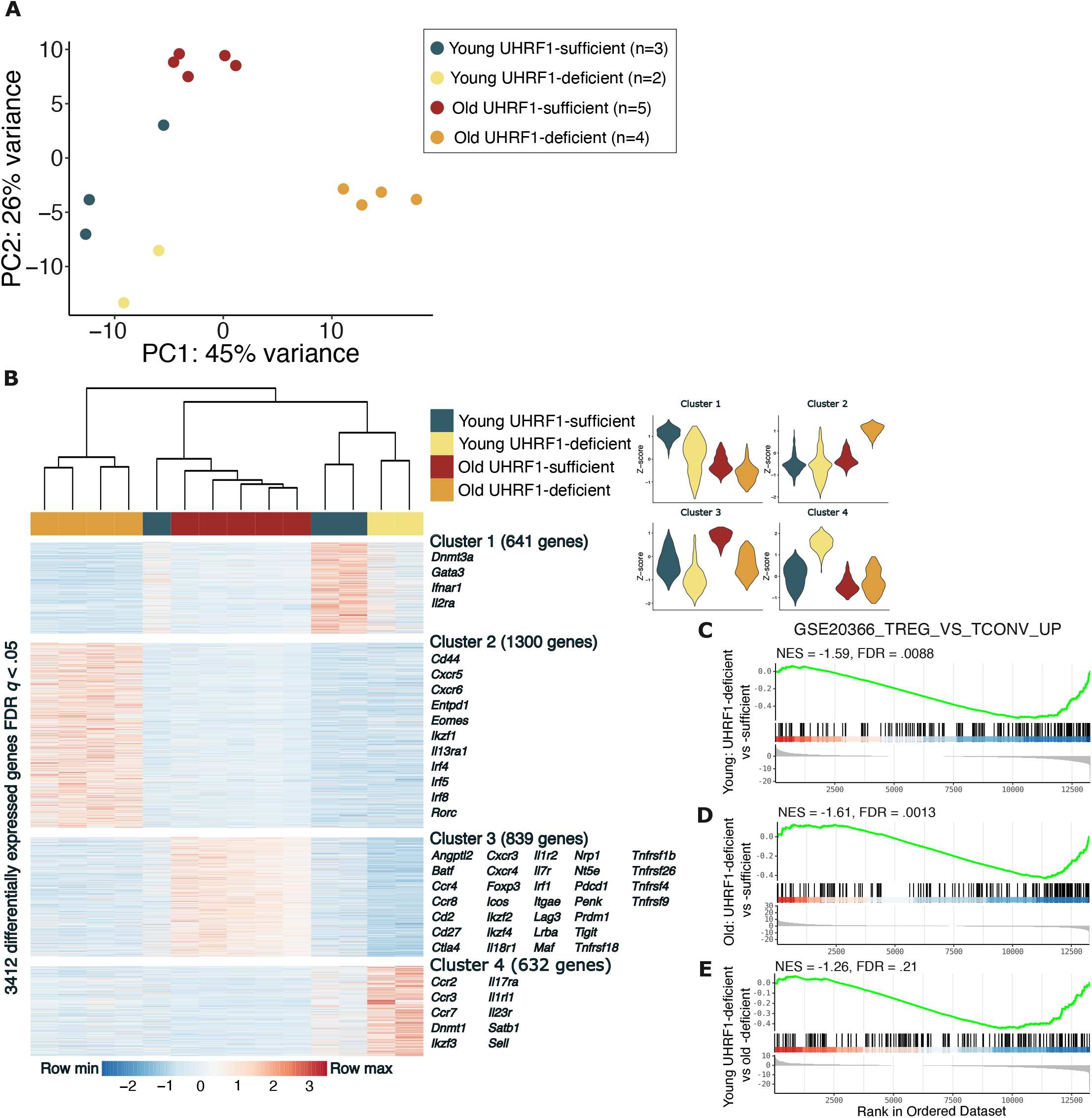
Maintenance DNA methylation is necessary for age-related alterations in Treg cell transcriptional signatures. **(A)** Principal component analysis (PCA) and **(B)** *K*-means clustering of 3,412 DEGs, identified from ANOVA-like testing with FDR *q* < 0.05 from Treg cells isolated from young and old UHRF1-chimeric mice. **(C)** Gene set enrichment analysis for regulatory (TREG) vs conventional T (TCONV) cell profiles between UHRF1-deficient vs -sufficient young mice; **(D)** UHRF1-deficient vs -sufficient old mice; and **(E)** young vs old UHRF1-deficient mice. *K*-means clustering used Ward.D2 clustering method. Violin plots show median and quartiles. Differential gene expression analysis *p*-adjusted cutoff = 0.05.

**Supplemental Figure 1.**
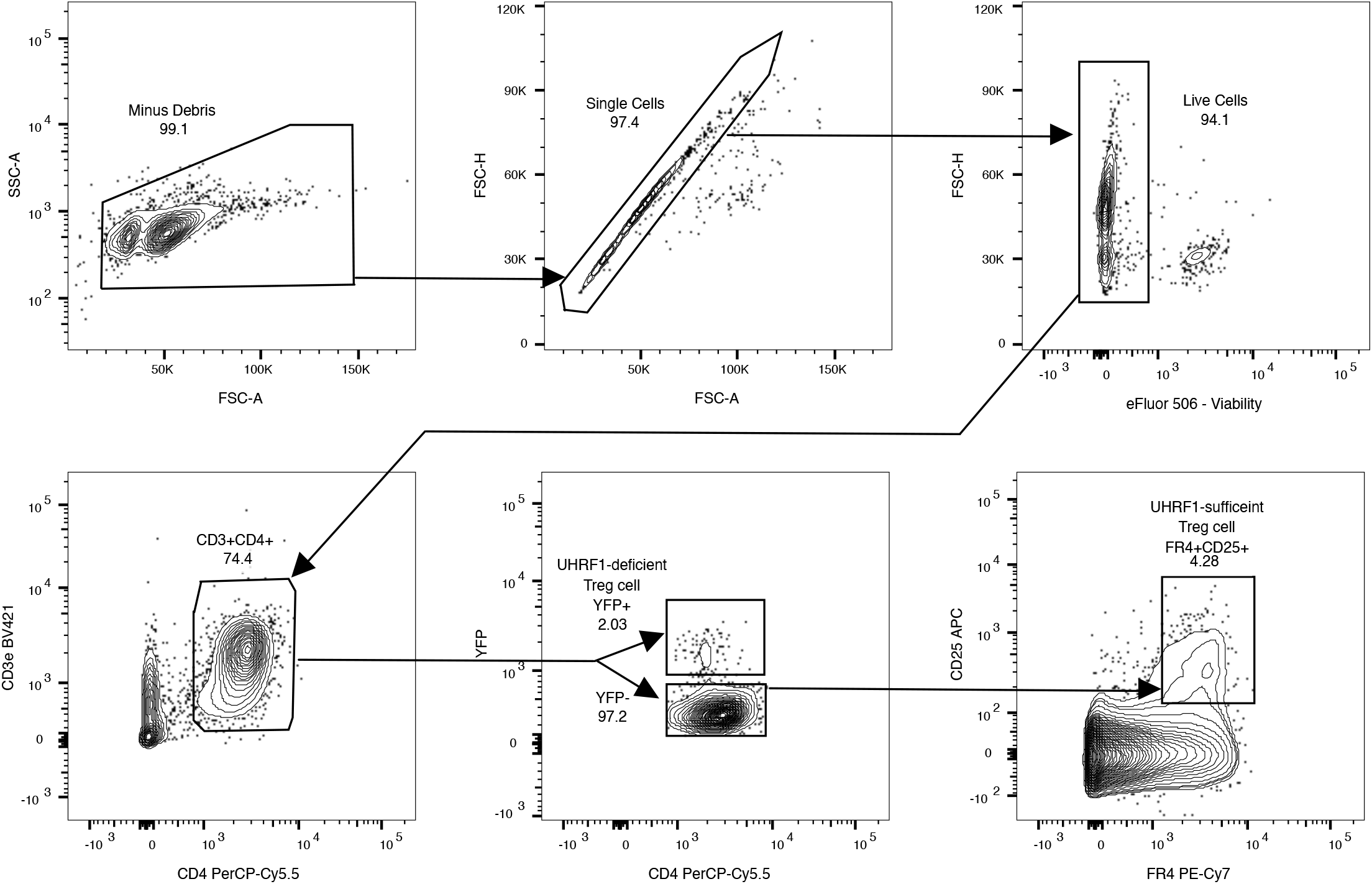
Sequential gating strategy used to sort live UHRF1-deficient CD3ε^+^CD4^+^Foxp3-YFP^+^ and UHRF1-sufficient CD3ε^+^CD4^+^Foxp3-YFP^-^CD25^+^FR4^+^ Treg cells for RNA-sequencing and mRRBS analysis.

Principal component analysis (PCA) revealed that 45 percent of the variance in the data set was due to differences between old UHRF1-deficient Treg cells and the other groups, with 26% of the variance due to differences between young UHRF1-deficient and old UHRF1-sufficent Treg cells **(Figure 1A)**, supporting the notion that UHRF1 is a determinant of transcriptional changes that occur in Treg cells over the lifespan. Unsupervised *K*-means clustering of young and old UHRF1-sufficient and -deficient Treg cells identified four clusters of 3,412 differentially expressed genes (FDR *q* < 0.05) (**Figure 1B**). Cluster 1 contained 641 differentially expressed genes upregulated in young UHRF1-sufficient Treg cells, including genes related to Treg cell lineage (*Gata3)* and function (*Il2ra, Ifnar1)*. Cluster 2, the largest group of differentially expressed genes, contained 1,300 genes upregulated in old UHRF1-deficient Treg cells, including genes related to Treg cell function and stability (*Cd44, Entpd1, Ikzf1)* as well as a variety of interferon regulatory factors (*Irf4, Irf5, Irf8)*. Cluster 3 contained 839 DEGs upregulated in old UHRF1-sufficient Treg cells and was comprised of genes canonically involved in Treg cell function, identity, and stability (*Foxp3, Ctla4, Lag3, Ikzf2, Icos, Il18r1, Nrp1*). Cluster 4 contained 632 differentially expressed genes upregulated in young UHRF1-deficient Treg cells, including genes involved in epigenetic regulation of Treg cell function (*Dnmt1, Satb1*) and identity (*Ikzf3*).

Consistent with loss of Treg cell identity upon disruption of DNA methylation patterning as previously described^9,10^, gene set enrichment analysis (GSEA) comparing young UHRF1-deficient and UHRF1-sufficient Treg cells revealed a shift from Treg to conventional T cell signatures (**Figure 1C**). We observed a similar pattern in the comparison between UHRF1-deficient and UHRF1-sufficient Treg cells in old mice (**Figure 1D)** as well as between young and old UHRF1-deficient Treg cells (**Figure 1E**).

Pairwise comparison between age-matched young and old UHRF1-deficient and -sufficient Treg cells, as well as between young UHRF1-deficient and old UHRF1-deficient Treg cells, demonstrated 655, 1,212, and 1,798 differentially expressed genes (FDR *q* < 0.05), respectively (**Supplemental Figure 2A-C**). Many of the shared genes downregulated in age-matched comparisons between UHRF1-deficient chimeric mice are canonically implicated in Treg cell activation and effector function, including *Pdcd1, Lag3, Icos, and Penk*, as well as chemokine receptor, *Ccr4*, which is required for Treg cell accumulation in mucosal tissues^14^, and *Ccr8*, which has been suggested as a therapeutic target on tumor Treg cells^15^ (**Supplemental Figure 2A-B**). Notably, Ikaros transcription factor family members necessary for Treg cell function, *Ikzf1, Ikzf2*, and *Ikzf4*^16–18^, were differentially expressed between old UHRF1-deficient and -sufficient Treg cells, with *Ikzf1* exhibiting upregulation in old UHRF1-deficient Treg cells (**Supplemental Figure 2B**). As previously reported, *Foxp3* was not differentially expressed between young UHRF1-deficient and -sufficient Treg cells^9^ but was downregulated in old UHRF1-deficient Treg cells (**Supplemental Figure 2A-B**). Direct comparison of young and old UHRF1-deficient Treg cells demonstrated that old UHRF1-deficient Treg cells downregulate core Treg cell genes such as *Foxp3, Ctla4, Il2ra, and Il1rl1*, and have higher expression of other genes associated with Treg cell function, such as *Entpd1, Lag3, Tnfrsf9*, and *Pdcd1* (**Supplemental Figure 2C**).

**Figure 2.**
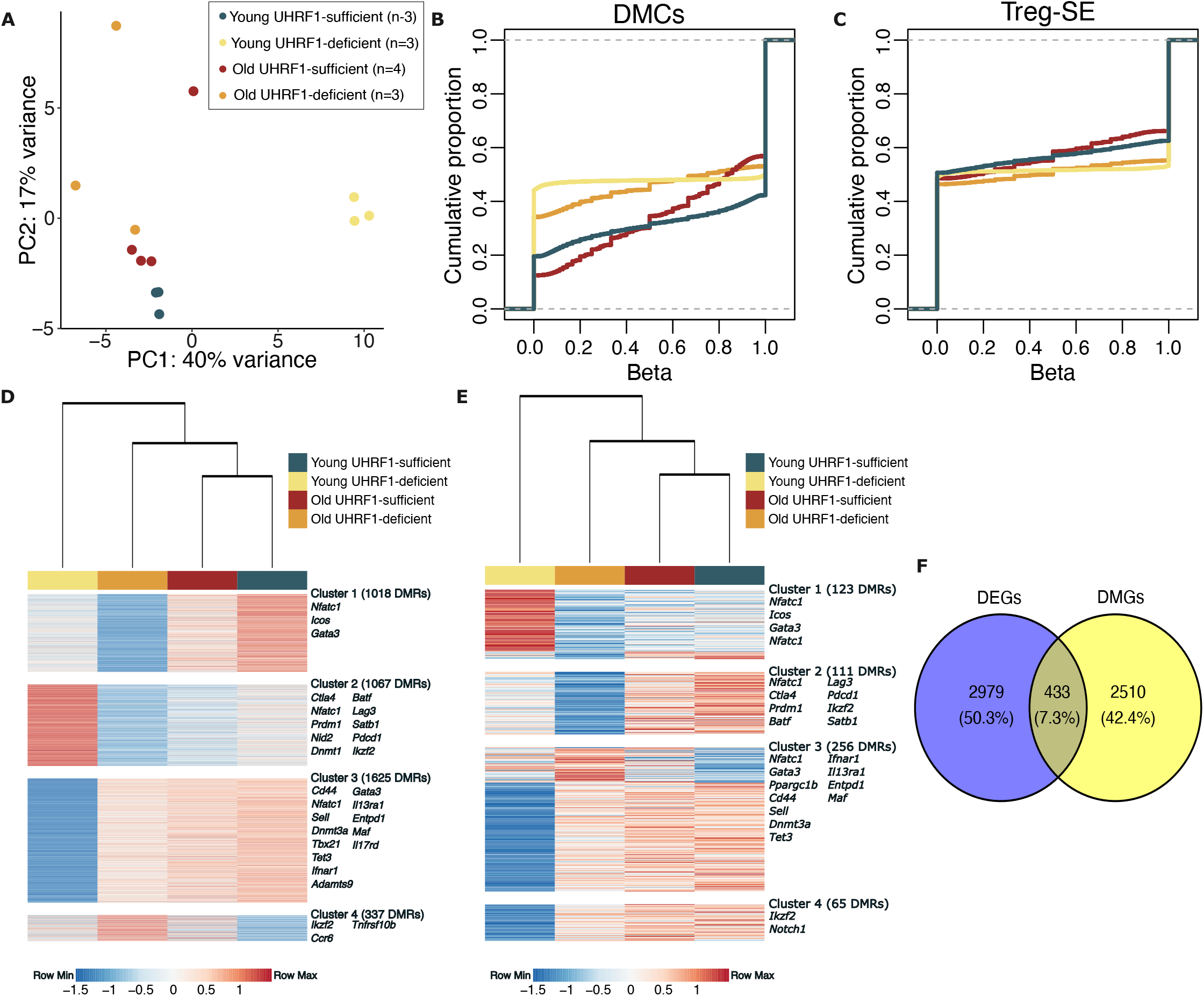
Maintenance DNA methylation is necessary for age-related alterations in Treg cell methylation signatures. **(A)** Principal component analysis (PCA) and **(B)** cumulative distribution plot of 110,561 differentially methylated cytosines (DMCs) between Treg cells isolated from young and old UHRF1-chimeric mice expressed as β scores, with 0 representing unmethylated and 1 representing fully methylated; a shift in the cumulative distribution function up and to the left represents relative hypomethylation. **(C)** Cumulative distribution plot of differentially methylated cytosines within the Treg super-enhancer (Treg-SE) regions. **(D)** *K*-means clustering of differentially methylated regions (DMRs) between Treg cells isolated from young and old UHRF1-chimeric mice. (**E)** *K*-means clustering of differentially methylated regions overlapping with differential gene expression from Figure 1B. **(F)** Venn diagram of differentially methylated genes (DMGs) and differentially expressed genes (DEGs). Differentially methylated cytosine analysis with *p*-adjusted cutoff = 0.05. Differentially methylated region analysis with *p* value < 0.05.

We then examined differences in gene expression between select genes implicated in Treg cell suppressive (**Supplemental Figure 2D**) and tissue effector (**Supplemental Figure 2E**) function alongside expression of transcription factors implicated in Treg cell function **(Supplemental Figure 2F**). Consistent with previous reports^8^ and movement toward increased effector Treg cells during aging^19^, old Treg cells had higher expression of many effector Treg cell genes (**Supplemental Figure 2E**). Furthermore, in old UHRF1-deficient Treg cells we noted downregulation of multiple genes in the TNFR superfamily **(Supplemental Figure 2G**), which encode key surface receptors that support Treg cell function and interaction with multiple immune cell subsets^20^ . Together, these data support that maintenance DNA methylation is required for Treg cell identity across the life course.

We used reduced representation bisulfite sequencing (RRBS) to determine whether the loss of UHRF1 affects age-related changes in DNA methylation. PCA of 110,561 differentially methylated CpGs (FDR *q* < 0.05) showed that 40 percent of the variance in the dataset was attributable to the differences between young UHRF1-deficient Treg cells and the other groups, while 17 percent of the variance was due to differences between young UHRF1-sufficient and old UHRF1-deficient Treg cells (**Figure 2A**). Consistent with previous results^9,10^, loss of UHRF1 led to a global hypomethylation pattern (**Figure 2B**), with a hypermethylation pattern within Treg super-enhancer regions (**Figure 2C**) in old and young Treg cells. An unsupervised procedure revealed 4,047 differentially methylated regions (DMRs) that were within 5 kb of 2,943 genes. *K-*means clustering of these DMRs revealed four clusters (**Figure 2D**). Notably, clusters 2 and 3 contained the largest number of differentially methylated regions, with each containing several genes canonically implicated in Treg cell function and stability in cluster 2 (e.g., *Ctla4, Prdm1, Lag3, Pdcd1, Ikzf2*). These DMRs were hypermethylated in young UHRF1-deficient Treg cells (cluster 2). Consistent with previous results^9^, DMRs within inflammatory genes, such as *Tbx21*, were hypomethylated in young UHRF1-deficient Treg cells but hypermethylated in old UHRF1-deficient Treg cells (cluster 3). Some DMRs overlapped the same gene, such as *Nfatc1*, with DMRs associated across multiple clusters. Of the 4,047 identified DMRs, 555 corresponded to genes that were also differentially expressed between the groups (**Figure 2E**), with 433 of them representing unique genes (**Figure 2F)**. Together, these data demonstrate that loss of UHRF1 results in alterations in DNA methylation patterns distinct with age.

**Supplemental Figure 2.**
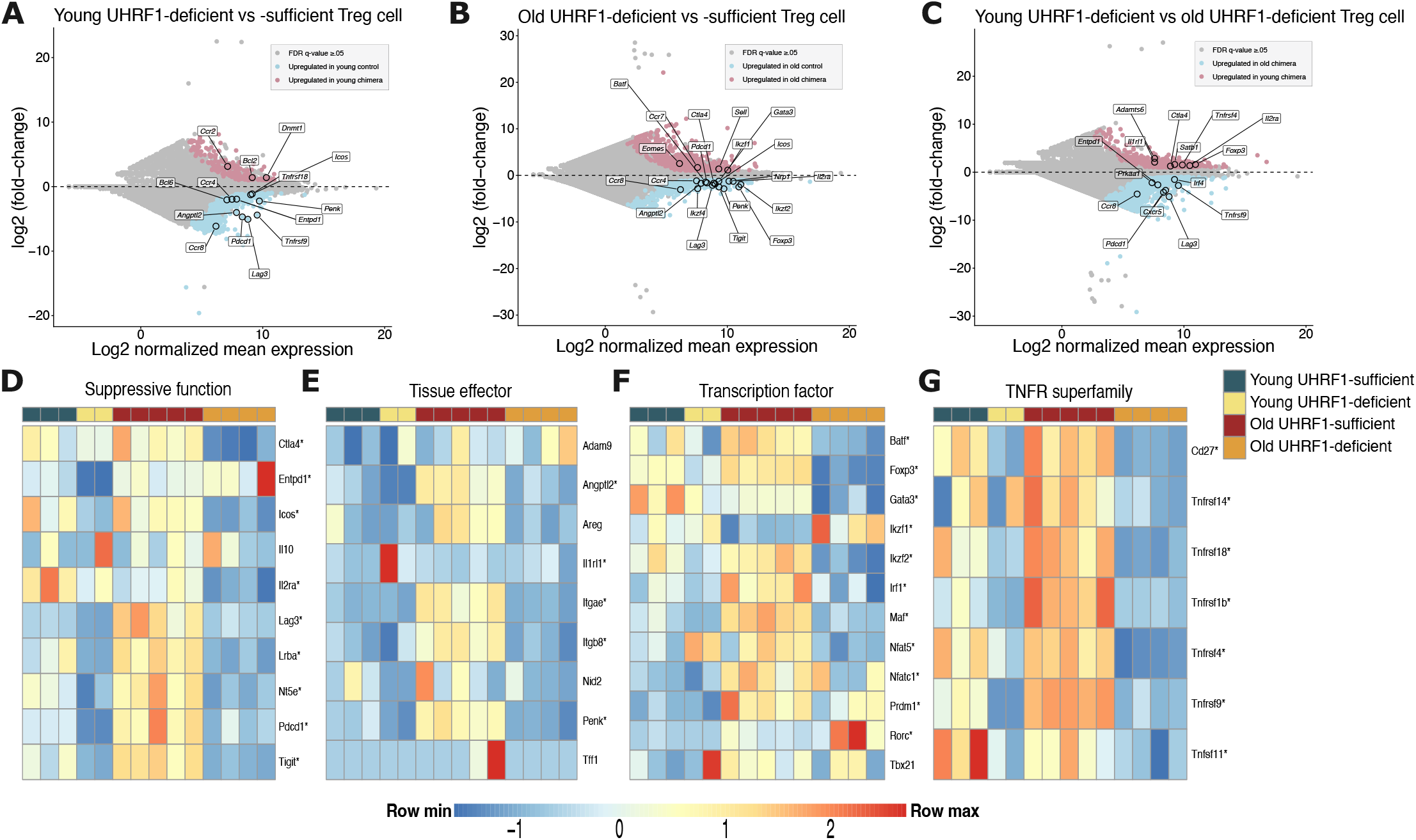
Loss of UHRF1 loss alters Treg cell transcriptional signatures and core gene expression with age. MA plot comparing gene expression between UHRF1-deficient CD3ε^+^CD4^+^YFP^+^ and control UHRF1-sufficient CD3ε^+^CD4^+^YFP^−^CD25^+^FR4^+^ splenic Treg cells from **(A)** young and **(B)** old UHRF1-chimeric mice. **(C)** MA plot comparing gene expression between young and old UHRF1-deficient splenic Treg cells. Expression of selected genes implicated in Treg cell **(D)** suppressive and **(E)** tissue effector function. **(F)** Expression of selected transcription factors canonically implicated in Treg cell function. **(G)** Expression of selected TNFR superfamily genes. * Denotes statistically significant differential expression from *K*-means clustering in Figure 1B.

## Discussion

Aging disrupts Treg cells’ requirement for a specific pattern of DNA methylation to establish cell identity as well as exert their suppressive and tissue-reparative functions^8–10^. Here, we demonstrated using young and old Treg cell-specific *Uhrf1*-chimeric mice that maintenance DNA methylation is necessary for age-related Treg cell transcriptional and DNA methylation signatures across the life course. Old Treg cells deficient in UHRF1 displayed a distinct transcriptional signature marked by higher expression of genes necessary for Treg cell function and stability (*Entpd1, Ikzf1, Cd44, Cxcr5*) while simultaneously downregulating other core Treg cell signature genes (*Foxp3, Ctla4, Il2ra, Nrp1, Ikzf2, Ikzf4*). We also observed age-dependent differences in DNA methylation between young and old UHRF1-deficient Treg cells. Young Treg cells displayed a hypomethylation phenotype around effector T cell inflammatory loci (**Figure 2D** Cluster 3, e.g. *Tbx21)* and a hypermethylation phenotype within genes implicated in Treg cell function and stability (**Figure 2D** Cluster 2, e.g. *Ctla4, Lag3, Dnmt1, Satb1*), consistent with prior work^9^. This pattern was not present in old UHRF1-deficient Treg cells, which displayed a phenotype more like young and old UHRF1-sufficient Treg cells, while still displaying a global hypomethylation profile and a hypermethylation profile at the Treg super-enhancer regions.

Given the central importance of DNA methylation in Treg cell function, numerous studies have sought to identify mechanisms that are necessary to maintain or promote the Treg cell-defining pattern of DNA methylation^21–25^. Cellular states that impede Treg cell function and are also linked to aging, such as impaired mitochondrial metabolism, altered Treg cell DNA methylation patterns^26–28^, and polymorphisms in Treg cell CpG-rich regions have been associated with human autoimmune disease^29^. While most of these studies were performed in young mice, during aging Treg cells experience cell-autonomous changes in their function across multiple tissues that contribute to dysfunctional repair of lung tissue after influenza pneumonia^8^ and impaired insulin sensitivity^30^.

For this study, we examined the differences between UHRF1-deficient and sufficient Treg cells within the same mouse by exploiting chimerism of Cre-recombinase active CD3ε^+^CD4^+^*Foxp3*-YFP^+^ (UHRF1-deficient) cells compared with CD3ε^+^CD4^+^*Foxp3*-YFP^−^CD25^hi^FR4^hi^ (UHRF1-sufficient) Treg cells, which we and others have previously validated^8,13^. Because the chimeric strategy requires two X chromosomes, all mice were obligate females. Whether our findings apply to males is unknown. As the UHRF1-sufficient population is sufficient to suppress lethal autoimmunity and permit normal aging^9^, this strategy allowed for control of any potential microenvironmental differences between mice that might impart a difference in our readouts. Nevertheless, the Treg cell pool is heterogeneous throughout lymphoid and non-lymphoid tissues^31,32^. Therefore, we are limited in our conclusions due to the use of bulk transcriptomic and DNA methylation analysis on splenic Treg cells. These results prompt future studies to examine how DNA methylation and factors that modulate DNA methylation and cell function differ between old and young Treg cells across tissue and inflammatory contexts.

## Materials and Methods

### Mice

All mice were housed and used in accordance with the Institutional Animal Care and Use Committee (IACUC) at Northwestern University. Animals received water ad libitum, were housed at a temperature range of 20°C–23°C under 14-hour light/10-hour dark cycles, and received standard rodent chow. C57BL/6 *Uhrf1*^fl/fl^ mice, which harbor loxP sequences flanking exon 4 (ENSMUSE00000139530) of the *Uhrf1* gene (ENSMUSG00000001228), were a gift from the laboratory of Srinivasan Yegnasubramanian (Johns Hopkins University, Baltimore, Maryland, USA). *Foxp3*^*YFP-Cre*^ mice were obtained from The Jackson Laboratory (stock no. 016959). As the *Foxp3-* driven Cre-recombinase is X-linked, Cre-mediated recombination is subject to random X-chromosome inactivation in females. Therefore, female mice heterozygous for the Cre allele and homozygous for the floxed *Uhrf1* allele (*Uhrf1*^*fl/fl*^*Foxp3*^*+/YFP-cre*^) were used and contained a mixed population of UHRF1-sufficient and -deficient Treg cells within the same animal^9^. As two copies of the X chromosome are necessary for this strategy, all mice are obligate females. Littermate controls were used for both young (3-month-old) and old mice (20-24-month-old) cohorts.

### Flow cytometry sorting

Single-cell suspensions were prepared from mouse spleens and enriched for CD4^+^T cells by positive selection using biotinylated anti-CD4 (clone GK1.5; Miltenyi, #130-102-024) and streptavidin-conjugated microbeads (Miltenyi, #130-048-1010) over magnetic columns. Enriched cells were stained with the antibody panel described in **Supplemental Table 1**, and dead cells excluded using a viability dye prior to sorting. Cell sorting was performed using the 4-way purity setting on BD FACSAria SORP instruments with FACSDiva software.

### RNA-sequencing, modified reduced representation bisulfite sequencing (mRRBS), and data analysis

For RNA-seq, flow cytometry-sorted cells (10^3^–10^5^) were immediately lysed in QIAGEN RLT Plus buffer containing 1% β-mercaptoethanol and processed for RNA and DNA extraction using the QIAGEN AllPrep Micro Kit, as described previously^9^. After sequencing, raw binary base call (BCL) files were converted to FASTQ files using bcl-convert 3.10.5 (Illumina). All FASTQ files were processed using the nf-core/RNA-seq pipeline version 3.9 implemented in Nextflow 23.04.2 with Northwestern University Quest HPC (Genomic Nodes) configuration (nextflow run nf-core/rnaseq - profile nu_genomics --genome GRCm38)^33–35^. In short, lane-level reads were trimmed using trimGalore! 0.6.5, aligned to the GRCm38 reference genome using STAR 2.7.11, and quantified using Salmon. After quantification, differential expression analysis was performed in R version 4.2.0 using DESeq2 v1.38.3.53. K-means clustering of differentially expressed genes (*q*<0.05) was performed using a previously published custom R function^36^. In brief, k was first determined using the elbow plot and the kmeans function in R stats 3.6.2 (Hartigan–Wong method with 25 random sets and a maximum of 1,000 iterations) was used for k-means clustering. Samples were clustered using Ward’s method and a heatmap was generated using pheatmap version 1.0.12.

For mRRBS, flow cytometry-sorted cells were first lysed with QIAGEN RLT Plus and then genomic DNA was extracted using the AllPrep DNA/RNA Micro Kit (QIAGEN). mRRBS DNA methylation libraries were generated as previously described^8–10,37^. In short, genomic DNA was fragmented with MspI (New England Biolabs), size-selected for 100–250-bp fragments using solid-phase reversible immobilization beads (MagBio Genomics), and then bisulfite converted with the EZ DNA Methylation-Lightning Kit (Zymo Research) per the manufacturer’s protocol^38^. Six libraries were prepared using the Pico Methyl-Seq Library Prep Kit (Zymo Research) and pooled in an equimolar ratio and sequenced using single-end reads (75 bp) with a NextSeq 2000 P2 reagent kit (100 Cycles; Illumina).

Methylation sequencing analysis was performed using our published Bismark-based pipelines^8–10,37,39^.

**Supplemental Table 1.** Flow cytometry reagents.

| Antigen | Conjugate | Company | Clone | Catalog No. |
| --- | --- | --- | --- | --- |
| CD3e | BV421 | eBioscience | 145-2C11 | 404-0031-82 |
| CD4 | PerCP-Cy5.5 | BioLegend | GK1.5 | 100434 |
| CD25 | APC | eBioscience | PC61.5 | 17-0251-82 |
| FR4 | PE-Cy7 | eBioscience | eBio12A5 | 25-5445-82 |
| Viability | eFluor 506 | eBioscience | N/A | 65-0866-14 |

## Data and material availability

Raw sequencing data will be made publicly available on the GEO database pending peer-reviewed publication.

## Author Contributions

JKG, QL, CPRF, and BDS contributed to the conception, hypothesis delineation, and design of the study. JKG, QL, CPRF, KAH, DHR, AMJ, BJU, HAV, EMS, and BDS performed experiments/data acquisition and analysis. JKG, QL, CPRF, and BDS wrote the manuscript.

## Acknowledgments

JKG is supported by NIH awards T32GM144295, T32HL076139, and F30AI188691. QL is supported by the David W. Cugell Fellowship and The Genomics Network (GeNe) Pilot Project Funding. CPRF is supported by NIH award T32HL076139. DHR is supported by NIH award T32GM144295 and the David W. Cugell Fellowship. AMJ is supported by NIH award F32HL162418 and the Parker B. Francis Opportunity Award. BDJ is supported by NIH awards R01HL149883, R01HL153122, P01HL154998, P01AG049665, U19AI135964, U19AI181102 and U54AG079754.

We wish to acknowledge the Northwestern University Flow Cytometry Core Facility supported by CA060553; the BD FACSAria SORP system was purchased with the support of S10OD011996. We also acknowledge the Northwestern University RNA-Seq Center/Genomics Lab of the Pulmonary and Critical Care Medicine and Rheumatology Divisions. This research was supported in part through the computational resources and staff contributions provided by the Genomics Compute Cluster, which is jointly supported by the Feinberg School of Medicine, the Center for Genetic Medicine, and Feinberg’s Department of Biochemistry and Molecular Genetics, the Office of the Provost, the Office for Research, and Northwestern Information Technology. The Genomics Compute Cluster is part of Quest, Northwestern University’s high-performance computing facility, with the purpose of advancing research in genomics.

